# The Gaussian Network Model as a Framework for Allosteric Analysis: Dynamic Distance, Edge Centrality, and Entropy Sensitivity in KRAS

**DOI:** 10.1101/2025.05.18.654696

**Authors:** Burak Erman

## Abstract

Allosteric communication in proteins relies on network connectivity patterns that channel conformational signals between distant sites. We introduce a unified mathematical framework based on three complementary measures of network organization derived from a single quantity. The first, the dynamic distance *R*_*ij*_, quantifies the mean-squared relative fluctuation between residue pairs. From this foundation, we derive two further metrics: the edge centrality, which identifies contacts critical for global connectivity by measuring their recurrence across all possible communication pathways, and the entropy sensitivity, which quantifies how perturbations to specific interactions alter system-wide flexibility. The mathematical structure shows that both topological centrality and thermodynamic sensitivity are linear functions of the dynamic distance. This derived unification demonstrates that residue pairs with high dynamic dissimilarity simultaneously function as flexible bottlenecks essential for allosteric communication. Applied to the oncoprotein KRAS, all three measures converge to identify the same residue pairs, corresponding to experimentally known allosteric sites. This convergence provides a unified graph-theoretical explanation for their functional importance. Analysis of the G12D mutation and adagrasib binding shows how local perturbations rewire global communication pathways, highlighting specific residue pairs that gain or lose importance as network bottlenecks.

## 1. Introduction

Proteins are dynamic, entropy-driven systems whose thermally induced fluctuations, coordinated across residues, enable critical biological functions including enzymatic activity, allostery, and ligand binding. Allosteric communication, the transmission of conformational signals between distant sites, relies on network-like connectivity patterns that channel information through specific pathways within the protein structure. Understanding these communication networks requires frameworks that can simultaneously capture local residue interactions and global connectivity patterns. Elastic Network Models, ENMs, provide a coarse-grained approach by representing proteins as networks of nodes, typically Cα atoms, connected by harmonic springs, enabling analysis of collective motions through normal mode analysis with minimal computational cost compared to atomistic molecular dynamics [1-5]. This network representation transforms the complex three-dimensional protein structure into a mathematical graph suitable for rigorous analysis using graph theory, where concepts like spanning trees, minimal connected subgraphs that preserve communication pathways, provide a framework for analyzing which residue interactions contribute to network connectivity patterns that may be relevant to allosteric communication.

Among ENMs, the Gaussian Network Model, GNM, stands out as a particularly simple approach that uses the full power of graph theory to decode protein dynamics. GNM maps the protein’s native structure onto a mathematical graph where nodes represent residues and edges denote harmonic interactions within a cutoff distance, typically 7-10 Å, [5]. This graph-theoretic representation transforms the complex three-dimensional protein structure into a topological network whose mathematical properties directly encode functional information. The resulting Kirchhoff Laplacian matrix serves as the mathematical core of the model, where nonzero off-diagonal entries denote connected residues and diagonal entries represent each residue’s connectivity degree [5]. Crucially, this matrix captures not merely the protein’s topology, but the fundamental graph-theoretic properties that govern information flow, community structure, and cooperative motions throughout the protein network.

The power of graph theory in GNM becomes evident through its eigenvalue decomposition of the Kirchhoff Laplacian matrix. The eigenvalues λ_i_ of this matrix encode the vibrational frequencies of collective modes, with each eigenvalue representing a distinct pattern of coordinated motion across the protein network. These eigenvalues form a spectrum that directly determines the configurational entropy of the system through the fundamental relationship 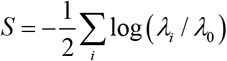 where *λ*_0_ is a reference constant and the sum excludes the trivial zero eigenvalue corresponding to global translation. This entropy-eigenvalue connection establishes a direct bridge between the graph-theoretic properties of the protein network and its thermodynamic behavior, which we will pursue in this study.

Graph theory, more specifically algebraic graph theory [6], provides GNM with tools to analyze protein networks beyond simple connectivity. Inverting the Kirchhoff matrix allows identification of indirect communication pathways between residues that are not directly connected, providing a quantitative view of how distant sites can influence each other through the network. Concepts such as betweenness centrality [7], clustering coefficients[8], and community detection algorithms [9, 10] can identify critical residues that serve as communication hubs, reveal densely connected functional modules, and map allosteric pathways through the protein network [11-15].

This study introduces a unified analytical framework based on three complementary measures of network organization. The first, dynamic distance, *R*_*ij*_, is a dimensionless metric that quantifies the mean-squared relative fluctuation between a pair of residues; increases in dynamic distance indicate that communication between the two residues relies on a larger number of alternative pathways through the network. The second is edge centrality, which weights dynamic distances by local topological connectivity and yields spanning tree probabilities that highlight edges critical for maintaining global communication. The third, entropy sensitivity quantifies how perturbations to specific interactions affect system-wide flexibility. While derived independently, all three measures are linear functions of *R*_*ij*_, indicating that changes in the number of alternative pathways, topological importance, and thermodynamic sensitivity share a common mathematical foundation in protein networks. Together, these three metrics provide a comprehensive framework for analyzing protein networks across multiple scales, from local residue couplings and structural connectivity to global dynamical sensitivity. This unified approach enables systematic characterization of the mechanisms underlying allosteric communication and conformational regulation in proteins.

When applied to KRAS, these three measures consistently identify the same set of residues, which correspond to experimentally known components of the allosteric hub. This convergence suggests that pathway diversity, topological importance, and thermodynamic sensitivity may reflect related aspects of protein network organization. The spectral properties of the Kirchhoff matrix, particularly the distribution of its eigenvalues, encode information about network modularity, bottlenecks, and the efficiency of information transfer between distant sites. Low-frequency modes (small eigenvalues) correspond to collective motions involving large, weakly connected regions, while high-frequency modes represent localized fluctuations in densely connected clusters [14, 15]. This spectral decomposition thus provides a natural hierarchy of motions, from global domain movements to local loop fluctuations, all encoded within the graph structure.

Unlike other ENMs, such as the Anisotropic Network Model (ANM) [16] that computes three-dimensional displacement vectors, GNM simplifies dynamics to isotropic, distance-based fluctuations. While this sacrifices directional information, it gains computational efficiency and, more importantly, enables the direct application of powerful graph-theoretic analysis tools. The scalar nature of GNM fluctuations allows for straightforward computation of residue cross-correlations from the inverse Kirchhoff matrix, showing the network’s inherent communication pathways and cooperative motion patterns that underlie allosteric regulation. These network-derived correlations have been successfully used to interpret allosteric regulation mechanisms in proteins [17-20].

GNM relies on several key assumptions that enable its graph-theoretic approach: (i) a multi-dimensional harmonic potential governs interactions, implying dynamics occur near the native state; (ii) residue fluctuations follow a Gaussian distribution, justified by the Central Limit Theorem due to many small thermal contributions; (iii) the equipartition theorem ensures equal energy distribution across vibrational modes; and (iv) interactions are limited to a cutoff distance, approximating local connectivity in the graph representation. These assumptions enable GNM to predict residue flexibility with high accuracy, correlating with experimental B-factors, while maintaining the mathematical precision necessary for graph-theoretic analysis.

While based on a simplified graph-theoretic representation, the GNM has demonstrated remarkable success in explaining diverse real-world protein phenomena through analysis of modified Kirchhoff matrices and their spectral properties. Notable applications include disease-related mutations [21-25], allosteric regulation and ligand binding [25, 26], and protein-protein interactions [27-32]. These applications demonstrate how GNM’s graph-theoretic approach, channeled through the spectral properties of the Kirchhoff matrix, provides functional insights that bridge the gap between protein structure and biological function, as discussed in detail below.

## 2. Theory

We represent proteins as networks, starting with the GNM to capture residue connectivity and motions, and then extending the analysis using graph-theoretic tools to explore network properties beyond the standard GNM. We use algebraic graph theory to provide a quantitative description of protein function, analyzing dynamic distance, spanning tree probabilities, and entropy sensitivity to show how residue flexibility, coordinated motions, and site-specific perturbations drive allosteric regulation and interactions.

A protein structure is mapped onto an undirected weighted graph with vertices *V* and edges *E*

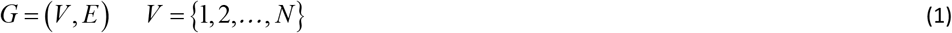

where, each vertex *i* ∈*V* represents the *C*_*α*_ atom of residue i, and an edge (*i, j*)∈ *E* is drawn if residues i and j are within a cutoff distance *r*_*c*_ in the equilibrium structure. Each edge carries a weight *w*_*ij*_ > 0 describing the strength of the interaction.

The weighted Laplacian (Kirchhoff) matrix *L* ∈ ℝ^*N*×*N*^ is defined by

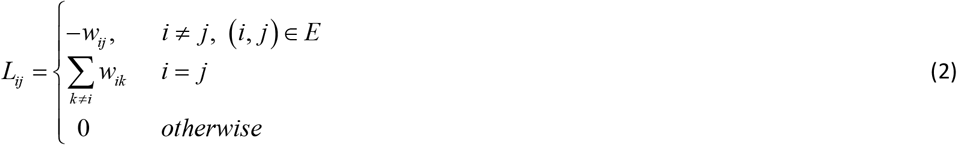

where, *w*_*ij*_ is the weight of an edge. Each diagonal element *L*_*ii*_ equals the weighted degree of vertex i. In GNM each *w*_*ij*_ is taken as unity. The Laplacian is often denoted as Γ in protein literature, but we retain the symbol *L* throughout this treatment.

### 2.1 Dynamic distance

Representing the instantaneus fluctuation Δ*R*_*i*_ of residue i as Δ*R*_*i*_ = *R*_*i*_ − ⟨*R*_*i*_⟩ where *R*_*i*_ is the position vector of residue i and the angular brackets is the time average, we introduce two complementary measures: the *Dynamic Distance, R*_*ij*_

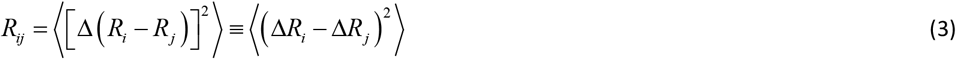

a scalar quantity that characterizes how rigidly or flexibly a pair of residues move relative to one another, with higher values indicating greater dissimilarity between their motions, and the normalized correlation coefficient *C*_*ij*_

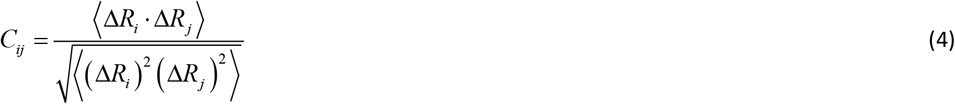

which quantifies whether their motions are aligned, opposed, or uncorrelated.

Dynamic distance *R*_*ij*_ quantifies relative motion (i.e., the dynamic dissimilarity) between residues i and j. In ENMs, it represents the mean-square fluctuation of the inter-residue distance, capturing how differently the residues move relative to each other. For connected residue pairs, this quantity relates to the probability that their edge participates in spanning trees, explained in more detail in the appendices A and C, providing a graph-theoretic interpretation of dynamic distance. When combined with the correlation coefficient *C*_*ij*_, dynamic distance helps characterize different types of residue relationships, distinguishing between coordinated motion, anticorrelated motion, and independent fluctuations. To fully characterize residue-residue dynamics, a single metric is insufficient. A large value of *R*_*ij*_ indicates that residues i and j undergo substantial relative motion, and examining *C*_*ij*_ simultaneously distinguishes whether this results from independent motion (*C*_*ij*_ ≈ 0) or coordinated but opposite motion (*C*_*ij*_ ≈ −1) . Similarly, while *C*_*ij*_ shows the directional relationship between displacements, *R*_*ij*_ provides the essential magnitude information. Together, they provide a more complete characterization of pairwise residue dynamics. We therefore use the pair (*R*_*ij*_, *C*_*ij*_) as a joint descriptor of residue motion relationships.

#### 2.1.1 Biological contexts for dynamic distance patterns

The dual-metric approach (*R*_*ij*_, *C*_*ij*_) distinguishes between different types of residue motion relationships:

1. Coordinated motion: Small *R*_*ij*_ with positive *C*_*ij*_ neighboring residues in secondary structures) indicates residues moving together (e.g.,
2. Independent motion: Large *R*_*ij*_ with *C*_*ij*_ ≈ 0 indicates uncoupled fluctuations (e.g., residues in different flexible loops)
3. Anticorrelated motion: Large *R*_*ij*_ with negative *C*_*ij*_ indicates residues moving in opposite directions (like regulatory switches, as their motions are anticorrelated with other parts of the protein)
4. Mixed flexibility: Large *R*_*ij*_ with *C*_*ij*_ ≈ 0 when one residue is flexible and another is rigid

The correlation coefficient *C*_*ij*_ provides directional information that *R*_*ij*_ alone cannot capture, enabling distinction between functionally different scenarios that produce similar dynamic distance values.

#### 2.1.2 Connection to the Gaussian Network Model (GNM)

In the GNM the time-averaged covariances of residue displacements are proportional to the Moore– Penrose pseudoinverse of the Kirchhoff (Laplacian) matrix

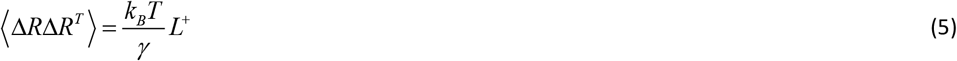

Here, *k*_*B*_ is the Boltzmann constant, *T* the temperature, *γ* is the single parameter of the GNM and *L*^+^ is the pseudoinverse of *L* . The ratio *k*_*B*_*T* / *γ* has dimensions of Length^2^ and *L*^+^ is dimensionless. In order to keep the dimensionless form of equations and to match GNM with standard graph theoretical terminology, we choose *k*_*B*_*T* / *γ* = 1 throughout the paper. With this choice the covariance equals the pseudoinverse of the Laplacian, 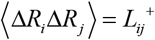.

Expanding Eq. 3 as

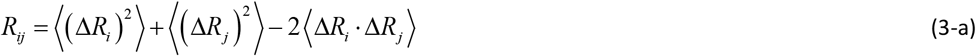

we obtain

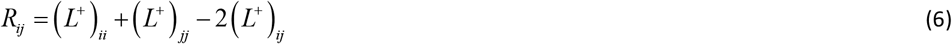

Dynamic distance *R*_*ij*_ is a non-negative scalar quantity (as will be shown in Section 2.4, for *w*_*ij*_ = 1, *R*_*ij*_ varies in the interval [0,1] under our dimensionless framework), and equation 6 sets the connection between *R*_*ij*_ and GNM. The weighted Kirchhoff/Laplacian *L* encodes local contacts. *L*^+^ encodes global connectivity and how thermal fluctuations and information propagate through the network.

The dynamic distance *R*_*ij*_ between residues i and j admits two equivalent interpretations. From the statistical-mechanical viewpoint of the Gaussian Network Model, it equals the mean-squared relative fluctuation 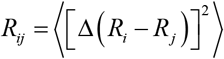 reflecting how strongly the motions of residues i and j are coordinated. From the graph-theoretic perspective, *R*_*ij*_ is determined by the Laplacian pseudoinverse and can be expressed through the Matrix Tree Theorem as the relative frequency with which communication pathways connecting i and j appear across all spanning trees of the network. Thus, the same quantity simultaneously measures relative motion in the fluctuation ensemble and pathway redundancy in the connectivity ensemble (See appendix C).

#### 2.1.3 Mode decomposition viewpoint

Writing the spectral decomposition of *L*,

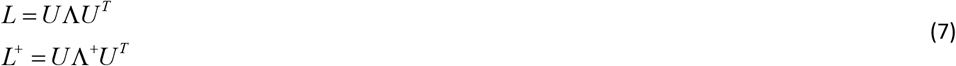

with ∧^+^ containing reciprocals of nonzero eigenvalues (zero for the null mode), we write

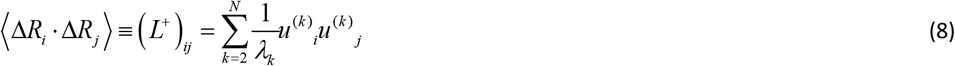

and

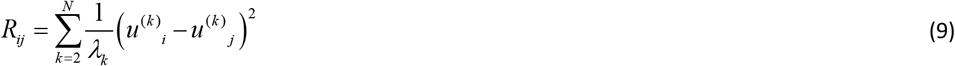

i.e. contributions from each nonzero (internal) mode k. This makes clear how global low-frequency modes dominate long-range dynamic distance.

Equation 9 may be written as the first term plus all the others as

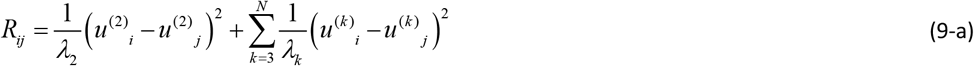

Each row i of *R*_*ij*_, when plotted as a function of j, represents the dynamic distance profile of residue i. Because the dominant mode typically provides the largest contribution, the dominant-mode approximation, 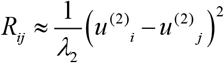, shows that all residues share the same peak positions in their profiles. Adding the remaining terms from Eq. 9-a modifies the amplitudes and may slightly adjusts the peaks, but the overall structure remains largely unchanged. Thus, analyzing a single residue’s dynamic distance profile is generally sufficient to approximate the essential pattern.

### 2.2 Edge centrality

A protein represented as a network can be studied at different levels: local residue-residue contacts, regional interactions such as domains, and global patterns of connectivity. To capture the latter in a minimal but comprehensive way, one needs a structure that connects all residues while avoiding redundancy. This role is fulfilled by a spanning tree: a subgraph that links all residues together with the smallest possible number of edges and no cycles [33].

Spanning trees therefore provide the simplest backbone that preserves communication across the <u>entire</u> protein. The total number of spanning trees quantifies the multiplicity of alternative ways in which all residues can be visited in the entire protein without forming cycles. By Kirchhoff’s classical matrix–tree theorem, this number can be calculated directly from the weighted Laplacian matrix of the graph [33].

By analyzing the ensemble of all possible spanning trees, one can assess which edges (residue-residue interactions) are essential for maintaining connectivity and which are replaceable. This insight forms the basis of the concept of edge centrality. The answer to the question ‘How often does a particular edge appear in all possible spanning trees?’ quantifies the importance of that edge in preserving connectivity.

The probability of a spanning tree T is given by the contribution of that tree’s edge weights divided by the sum of the contributions of all spanning trees:

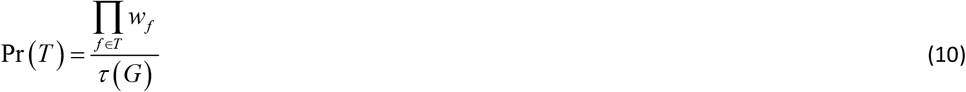

where the numerator is the product of the edge weights in T, and the denominator τ(G) is the sum of such products over all spanning trees. This expression defines a probability distribution over weighted spanning trees. The probability that a given edge e appears in a spanning tree drawn from this distribution is then

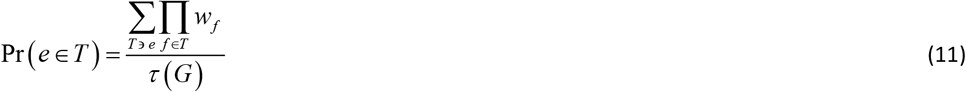

Here, the numerator is the sum of the weights of all spanning trees that contain e, and the denominator τ(G) is again the sum of such weights over all spanning trees. This gives a normalized probability between 0 and 1. Using Kircchoff’s matrix tree theorem [33] (See Appendix A) we show that this probability is equal to *w*_*ij*_ *R*_*ij*_ . We refer to this measure as the centrality of the edge. Thus, we define the spanning-tree edge centrality of an edge (i,j) as the fraction of spanning trees that include it

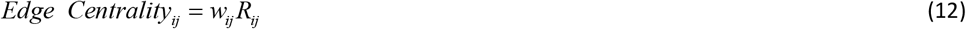

This quantity, wich varies in the interval [0,1], provides a direct and natural measure of an edge’s role in allosteric communication. Although edges are inherently local objects, their recurrence across the ensemble of spanning trees reflects their contribution to nonlocal connectivity: an edge that recurs across many spanning trees ensures that distant residues remain dynamically linked, thereby shaping the global communication backbone of the protein. In this sense, spanning-tree edge centrality bridges local contacts with system-wide communication pathways, offering a graph-theoretic perspective on how subtle perturbations at individual residues can propagate through the network to drive allosteric activity.

We note an important distinction in terminology: Edge centrality (*w*_*ij*_ *R*_*ij*_) applies only to connected pairs where *w*_*ij*_ > 0, while dynamic distance *R*_*ij*_ provides information about both local (connected) and long-range (distant) residue relationships.

### 2.3 Entropy response to edge perturbation. Entropy sensitivity

In this section, we evaluate the change in entropy of the system when an edge is perturbed. This is a global response to a local change in the protein.

The entropy of the graph is [15]

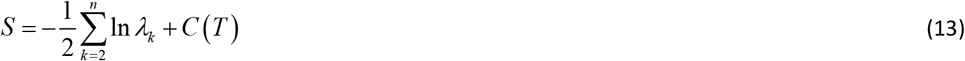

where, *C* (*T*) is a constant that depends on temperature. We pick one edge (i,j) and perturb its weight:

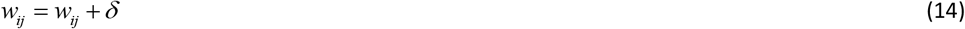

Then

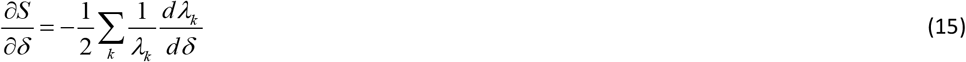

and using the standard Rayleigh–Schrödinger (first-order) perturbation (Explained in Appendix B) we obtain

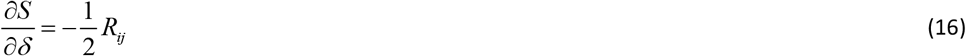

The negative sign says that increasing the spring/edge strength *w*_*ij*_ (adding δ>0) reduces the entropy, stiffening reduces available configurational phase space. Thus, this equation relates a local quantity to a global one.

Since ∂*S* / ∂*δ* varies in the interval [-0.5,0], strengthening an interaction(*δ* > 0) always decreases entropy. To better represent an edge’s capacity to increase flexibility, Figure 5 below instead plots the resulting entropy increase for a unit weakening of the edge, calculated as 0.5*R*_*ij*_ ∈^+^0, 0.5^+^ .

### 2.5 The unifying metric *R*_*ij*_

In summary, our analysis demonstrates that dynamic distance, edge centrality, and entropy sensitivity are all linear functions of the fundamental quantity *R*_*ij*_ . While dynamic distance captures the dynamic dissimilarity between residues, edge centrality reflects an edge’s importance in maintaining communication pathways, and entropy sensitivity quantifies the influence of each interaction on global network flexibility, all three metrics share a common foundation. This unified framework highlights how diverse aspects of residue-residue interactions, mechanistic coupling, network connectivity, and system-wide influence, can be consistently quantified using *R*_*ij*_, providing a coherent approach for analyzing allosteric communication in proteins.

Mutations and ligand binding perturb the dynamic distance between residues, quantified by the dual metric (*R*_*ij*_, *C*_*ij*_), thereby reshaping the protein’s communication network through interconnected local and global effects. These perturbations simultaneously operate across scales: locally, they weaken, strengthen, or reorganize specific dynamic distance, shifting the balance of fluctuation amplitudes and correlations; globally, they modify the ensemble of spanning trees and alter the entropy. The direct connection between local and global scales is captured by changes in the three centralities defined here. Perturbations of locally critical couplings propagate to produce disproportionate global effects on network adaptability, while global changes in entropy reshape the local landscape of residue interactions.

## 3. Application to KRAS

In this section, we apply the present theoretical framework to KRAS. We set all *w*_*ij*_ = 1 for all contacts, equivalent to the Gaussian Network Model, and analyze how dynamic distance, spanning-tree edge centrality, and entropy sensitivity respond to mutation and ligand binding, showing how communication pathways are rewired.

KRAS is a small GTPase that functions as a molecular switch, cycling between an active GTP-bound and inactive GDP-bound state to regulate critical cellular processes, including proliferation, survival, and differentiation. Its activity is tightly controlled by upstream signals from growth factor receptors (e.g., EGFR) and modulated by GEFs (guanine nucleotide exchange factors) and GAPs (GTPase-activating proteins). Upon activation, KRAS interacts with downstream effectors, primarily through the switch I (residues 30–38, including the critical T35 and D38) and switch II (residues 60–76, featuring G60 and Q61) regions, which undergo conformational changes upon GTP binding to recruit RAF, PI3K, and RALGDS. Oncogenic mutations (e.g., G12V, G12D, G13D, Q61H) impair GTP hydrolysis, locking KRAS in a constitutively active state and driving uncontrolled signaling through the MAPK/ERK and PI3K/AKT pathways, resulting in various types of cancers including lung and pancreatic cancers.

The allosteric network of KRAS involves long-range communication between the GTP-binding site (P-loop, residues 10–16), switch regions, and helix α3 (residues 92–110). Key residues like Y71, R73, and E107 form an allosteric hub that couples nucleotide state to effector binding. Membrane localization is mediated by the C-terminal hypervariable region (HVR, residues 166–172), which plays role in anchoring KRAS to the plasma membrane. This membrane association is critical for signal propagation, as it positions KRAS near upstream activators (e.g., receptor tyrosine kinases) and facilitates the spatial organization of downstream signaling complexes. The orientation of helix α4 (residues 151–165) and the membrane-proximal linker region further regulate the accessibility of switch regions, ensuring efficient signal relay from the membrane to the effector-binding domains [34].

The allosteric and dynamic properties of KRAS have been extensively studied and the critical residues are well identified. In particular, single-molecule FRET and NMR experiments have shown allosteric states and identified residues involved in allostery. These findings will be discussed after the KRAS analysis section and compared with the outcome of the theory.

### 3.1 Edge centrality

As discussed in the theory section, edge centrality quantifies how often a specific residue-residue contact contributes to maintaining the global connectivity of the structure. Edges with high centrality are those that repeatedly appear in spanning trees, and are indispensable for linking distant regions of the protein. As a result, such contacts serve as ‘communication highways’ through which long-range interactions are funneled. Values of edge centrality *w*_*ij*_ *R*_*ij*_ are presented in the left panel of Figure 1 for wild type KRAS using the Protein Data Bank structure 6GOD. The GNM is adopted where *w*_*ij*_ are taken as 1 for all contacting residues and 0 otherwise. The cutoff distance is taken as 7.8 Å in forming the Kircchoff, i.e, Laplacian matrix according to Eq 2. Values of *R*_*ij*_ are calculated from Eqs. 3-a and Eq. 6.

**Figure 1.**
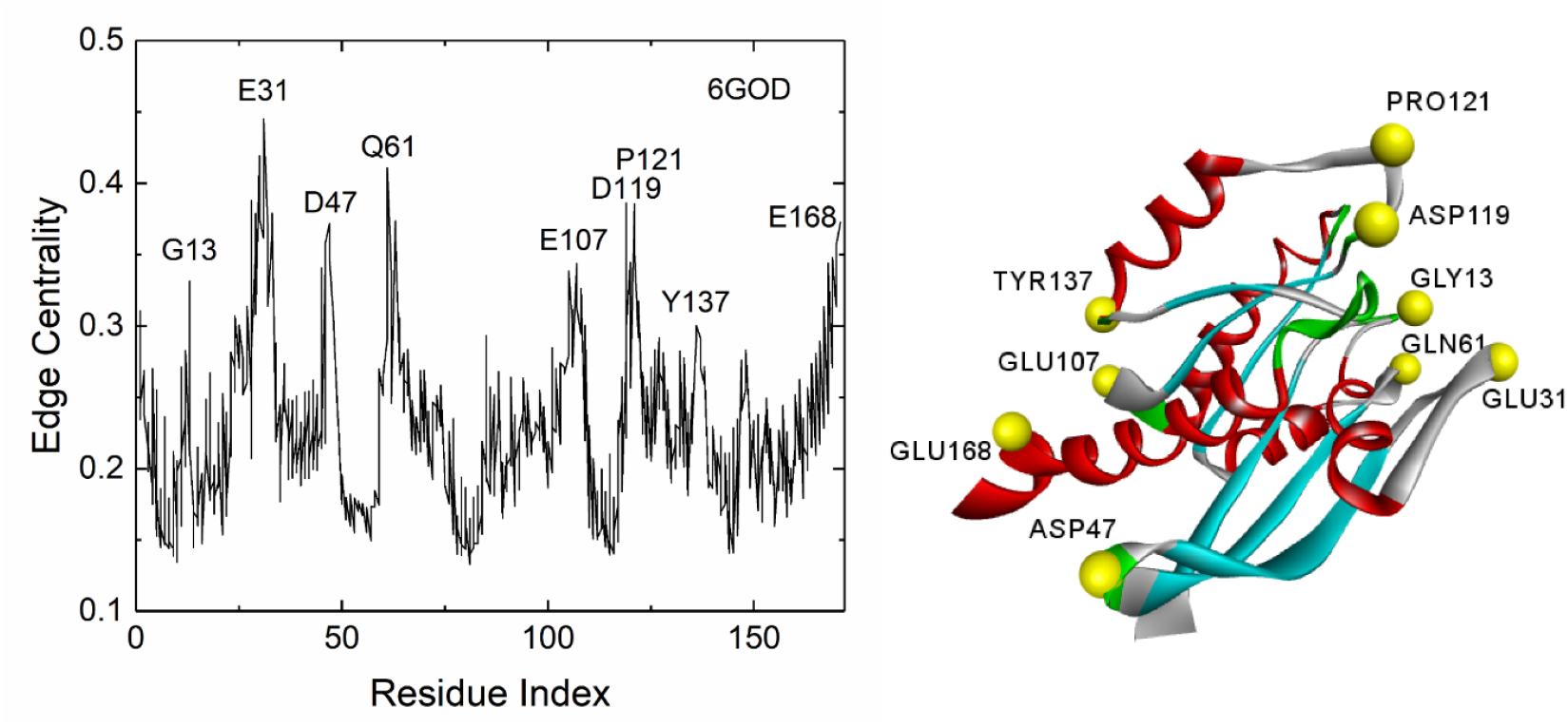
Left panel: Edge centrality profile of wild-type KRAS, 6GOD.pdb. For each residue i (abscissa), the ordinate shows the *R*_*ij*_ values with all contacting residues j. Only the off-diagonal elements of *R*_*ij*_ are used. Here, all edges incident on each residue i are displayed, and the scatter of values shows root-mean-squared deviations ranging from 0.088 to 0.122. Residues at the peak values, symbolically representing the ‘communication highway,’ are labeled. Right panel: Spatial locations of residues with peak edge centralities are mapped onto the three-dimensional structure of the protein. Calculations were performed using the Gaussian Network Model (GNM) with unit weights for all edge weights, *w*_*ij*_, and a cutoff radius of 7.8 Å. Ordinate values correspond to the fraction of spanning trees containing the given edge, ranging from 0 to 1.

All peak residues indicated in Figure 1 participate in the allosteric activity of KRAS. The most prominent peak corresponds to residue E31, which forms edges with V29, D30, Y32, and D33. Among these, the D30–E31 edge is present in 42% of all possible spanning trees that can be drawn on KRAS, corresponding to an edge probability of 0.42.

The right panel in Figure 1 shows that residues with high edge centrality are located on the protein surface. Surface residues often display high edge centrality because they have fewer possible contacts than buried residues. When spanning trees are drawn, these limited contacts are selected repeatedly as the only ways to connect surface residues to the rest of the protein. In network terms, such edges act as bottlenecks, communication gateways, funneling information flow through narrow channels along the ‘communication highway.’ Biologically, this means that surface residues are positioned to mediate allosteric communication, as signals traveling through the protein must frequently pass through these gateway contacts. In the extreme case where an edge ij provides the only connection of a residue to the network, all spanning trees must include ij, giving it a probability of 1.0. Thus, the higher the edge centrality value, the higher the probability that the edge lies on the main communication pathway.

In this and the following sections, the cutoff radius *r*_*c*_, which defines the extent of the first coordination shell around each residue, is chosen as 7.8 Å. In the literature, *r*_*c*_ is generally chosen in the range 6–10 Å for Cα representations, since this interval captures the dominant noncovalent contacts without over-densifying the network. The value we chose falls in the middle of the empirically established interval and has been widely used in previous ENM/GNM studies to balance inclusion of relevant side-chain packing interactions with computational sparsity of the Kirchhoff matrix. To assess sensitivity of our calculated results, we evaluated the derivatives of the mean dynamic distance ⟨*R*_*ij*_ ⟩ over all residue pairs of KRAS. For *r*_*c*_ ≥ 7.8 Å, the second derivative was essentially zero, indicating that the curve enters a linear regime. Linear regression of the first derivative over this range yielded a slope of 0.013±0.034 (95% CI), consistent with no systematic trend. Thus, beyond 7.8 Å, ⟨*R*_*ij*_⟩ decreases smoothly at a nearly constant rate, and the qualitative patterns reported in this work are insensitive to the precise cutoff.

The switch 1 edge E31-D30 contains 42% of all possible spanning trees and the fluctuations of E31 and D30 have correlation *C*_30,31_ of 0.48. The switch 2 residue Q61 makes an edge with G12 with a probability of 33% and a correlation of 0.26. The edges of D119 to P121 participate in nucleotide stability and downstream communication. The switch 2 residue Q61 has the next highest edge centrality edges. The residue D47 is located in the interswitch loop 3 region, which is involved in allosteric pathways across the Ras dimer interface [35], and D47 and E49 make stable salt bridge interactions with helix 5 residues R161 and R164 [36]. E107 is located in the allosteric site, a charged pocket formed at the interface of helix 3, loop 7, and helix 4. These interactions control the transitions between active and inactive conformations, influencing the protein’s allosteric regulation and signaling function [37]. Another group of peaks at D116 to P119 lies within the terminal part of the catalytic domain and contribute to the drug binding and allosteric regulation capabilities of KRAS [38]. Y137 plays a critical role in the allosteric regulation of the protein by forming a hydrogen bond with H94 in helix 3, which stabilizes the R-state conformation involving helix 3 and helix 4 interactions [39]. Finally, E168 has a critical role on dimerization through allosteric interactions [40].

### 3.2 Network reorganization upon mutation and ligand binding

Mutations and ligand binding perturb the protein’s dynamic landscape, redistributing how thermal fluctuations explore configurational space and reorganizing the pathways through which allosteric signals propagate. From the unified perspective developed in the theory section, changes in dynamic distance *R*_*ij*_ reflect alterations in both motion coordination and network topology: when *R*_*ij*_ increases for a residue pair, that interaction becomes a more critical bottleneck through which spanning trees must pass, thereby expanding the accessible configurational phase space, the volume of possible relative configurations the system can explore while maintaining network connectivity. This indicates enhanced dependence on that specific connection for maintaining network-wide communication; Conversely, when *R*_*ij*_ decreases, alternative edges with larger dynamic distances become the preferred routes, reducing the network’s reliance on the given i,j connection. These changes do not simply strengthen or weaken individual interactions, they fundamentally reroute communication pathways through the allosteric network. The functional consequences of such reorganization depend on biological context. In wild-type protein, critical bottlenecks (high *R*_*ij*_ edges) are necessary for efficient allosteric signaling, serving as key communication nodes that enable coordinated responses to regulatory inputs. For therapeutic inhibitors, creating redundancy by decreasing *R*_*ij*_ at these critical edges is desirable, as it disrupts the communication pathways essential for abnormal signaling. For oncogenic mutations, if the mutation increases *R*_*ij*_ at specific edges, it may establish new critical pathways that enhance constitutive signaling and drive uncontrolled cellular responses. In the following sections, we analyze how the G12D mutation and adagrasib binding reshape dynamic distance patterns in KRAS, identifying which residue pairs become more critical as communication bottlenecks and which become redundant as alternative pathways emerge.

#### 3.2.1 Dynamic distance profiles and changes upon G12D mutation

For a given reference residue i, the dynamic distance profile *R*_*ij*_ plotted against j shows which residues move most differently from the reference. Because the smallest nonzero eigenvalue and its corresponding eigenvector dominate the spectral decomposition (Eq. 9-a), peak positions in dynamic distance profiles are approximately independent of which residue is chosen as reference. However, the peak amplitudes depend on the choice of reference residue, as demonstrated in the figures below.

Figure 2 presents dynamic distance profiles for wild-type KRAS with G12 as reference (left panel) and with E31 as reference (right panel), calculated using Eqs3-a and 6 for GNM with r_c_=7.8 Å. Peak residues are labeled in each profile. The analysis identifies a set of residue pairs that consistently show high dynamic distances across different reference choices, corresponding to known components of the allosteric hub in KRAS. In agreement with the validity of Eq. 9-a and the discussion theren, while peak positions are conserved, the amplitudes vary: E31 exhibits larger dynamic distances than G12, indicating it is more dynamically separated from the protein network.

**Figure 2.**
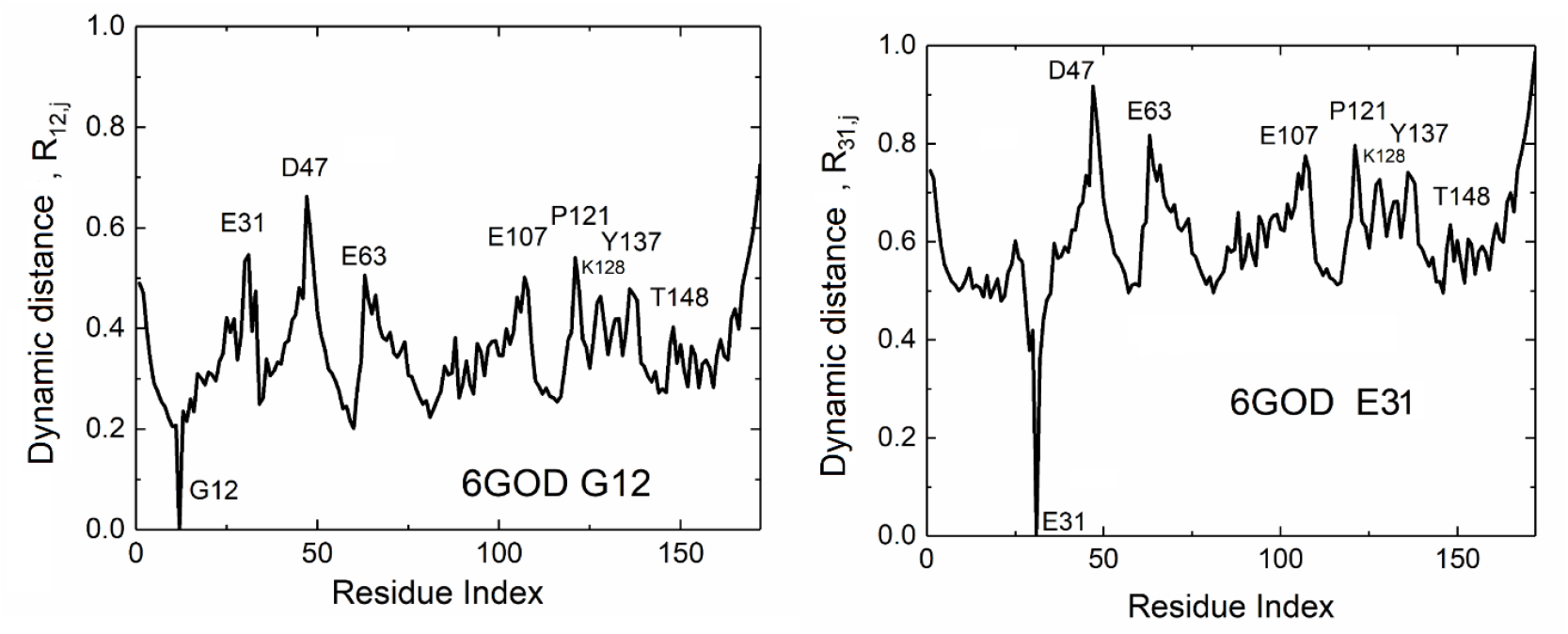
Dynamic distance profiles for G12 (left panel) and E31 (right panel) obtained for the wild type KRAS (6GOD.pdb) using GNM with a cutoff value of 7.8 Å. Residues of dominant peak values are identified. Same limits for the ordinates are used to facilitate easy comparison. Since *R*_*ii*_ = 0 by definition, the ordinate values for the reference residue is zero. Values of *R*_*ij*_ are calculated from Eqs. 3-a and Eq. 6

Calculations of dynamic distance profiles for the mutant, 6GOF.pdb, showed that peak positions remain identical to those of the wild type but the amplitudes show substantial changes relative to those of the wild type. Figure 3 presents the difference *R*_31, *j*_ (*G*12*D*) − *R*_31, *j*_ (*WT*) for E31 as the reference residue. Since absolute *R*_*ij*_ values depend on the overall network structure and eigenvalue spectrum, we converted dynamic distances to z-scores within each structure before calculating differences: Δ*z*_*ij*_ = *z*_*ij*_ (*G*12*D*) − *z*_*ij*_ (*WT*) . This normalization removes dependence on absolute scale differences while preserving relative pattern changes.

**Figure 3.**
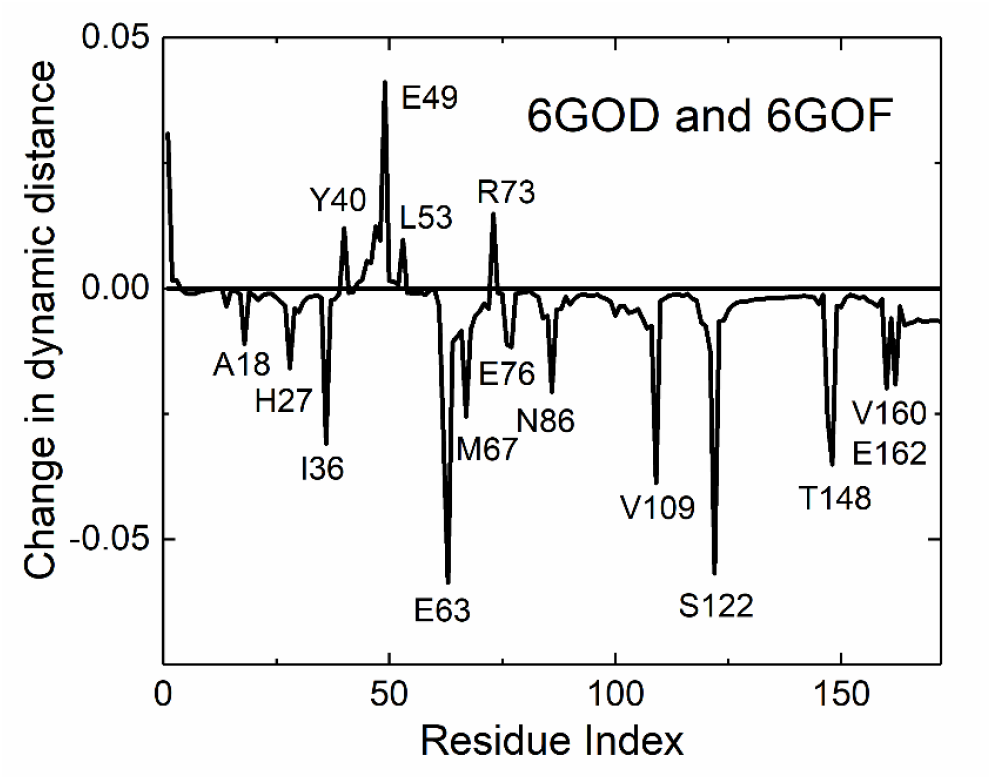
Change in dynamic distance ΔR_31_,_j_ upon G12D mutation. The ordinate shows the difference in dynamic distance between E31 and all other residues j between mutant (6GOF.pdb) and wild-type KRAS (6GOD.pdb). E31 was selected as the reference residue because it exhibits one of the highest edge centralities in switch 1 (Figure 1). Positive values indicate increased spanning tree paths pssing through E31 and the residue of the peak; negative values indicate decreased spanning tree path. The peak positions are approximately conserved when other high-centrality residues are used as reference points, consistent with the dominant eigenmode analysis (Eq. 9-a). Values of *R*_*ij*_ are calculated from Eqs. 3-a and Eq. 6

Positive peaks indicate residue pairs whose dynamic distance increases upon mutation, reflecting a rewiring of the communication network that enhances the number of available pathways connecting them, while negative peaks indicate residue pairs whose dynamic distance decreases, showing that their connection has become less critical as alternative edges become more important. The G12D mutation increases dynamic distances to Y40, E49, L53, and R73. The largest increase occurs for E49. A closer look at the interactions of E49 shows that its largest interactions are with its spatially neighboring helix residues. Although E49 and K172 are spatially distant, 86% of spanning trees in the WT structure pass through both residues, increasing to 90% in the G12D variant, highlighting how the mutation reshapes communication pathways and reinforces this long-range conduit. Similar increases are observed for other local and nonlocal residue pairs upon mutation. The increased dynamic distance of E49 upon G12D mutation provides a mechanistic insight into how this residue participates in long-range allosteric networks connecting the mutation site to effector-binding regions. NMR studies have identified E49 as part of KRAS dimerization surfaces, where it contributes to alternative dimer conformations [41]. Clinical sequencing has detected the E49K variant in patient samples with oncogenic potential [42]. Structural analyses of Ras-Raf complexes show that the D47-E49 region contacts the Raf-CRD/α5 domain, implicating this region in allosteric coupling [43]. Systematic mutagenesis of the α4-α5 region (including E49) demonstrates context-dependent effects on both dimerization and downstream signaling [35, 44, 45]. While these studies support the functional relevance of E49, the correspondence between the mechanistic observations and their impact on KRAS function remains to be fully established.

The interactions of E63, a switch 2 residue, show significant decrease upon mutation. Although E63 and K172 are separated by ca. 40 Å, 96% of spanning trees passing through both residues, decreases to 88% in the G12D variant. While *R*_*ij*_ decreases for the E63–K172 pair upon mutation, their fluctuations remain anticorrelated at the same level. This suggests that the mutation weakens their dynamic coupling yet preserves the mode of their relative motion, while rewiring global network pathways. Experimentally, cellular assays showed that the G12D and E63K mutants both lead to increased cell proliferation and activation of downstream effectors, indicating that although G12 and E63 occupy distinct structural roles, mutations at these sites can produce comparable functional effects on KRAS allostery [46].

#### 3.2.2 Dynamic distance changes upon ligand binding

Ligand binding alters dynamic distance patterns by modifying how residue fluctuations are coordinated across the protein network. To compare GDP-bound KRAS with and without inhibitor, we analyzed wild-type GDP-KRAS (6MBU.pdb) and adagrasib-bound GDP-KRAS (9O0R.pdb). Since absolute *R*_*ij*_ values depend on the overall network structure and eigenvalue spectrum, we converted dynamic distances to z-scores within each structure before calculating differences: Δ*z*_*ij*_ = *z*_*ij*_ (*ligand bound*) − *z*_*ij*_ (*apo*) . This normalization removes dependence on absolute scale differences while preserving relative pattern changes.

Figure 4 shows the Δ*z*_31, *j*_ profile with E31 as the reference residue. The peak positions remain consistent when other reference residues are used, confirming these patterns reflect genuine structural reorganization rather than reference-dependent artifacts. Adagrasib binds to KRAS at G13 to A18, F28, D30, N116, K117, D119, L120, S145, A146, K147. Among these, D108 and T148 are distant from the ligand-binding sites. Upon ligand binding, the fraction of spanning trees containing both residues decreases from 61% to 52%, reducing their contribution to global communication. At the same time, their fluctuations remain anticorrelated, with the correlation coefficient shifting slightly from -0.05 to -0.08, suggesting a strengthening of their out-of-phase motions despite the overall reduction in communication capacity.They are known to contribute to membrane orientation and signaling complex formation and are assigned as potential anticancer targets [47-49].

**Figure 4.**
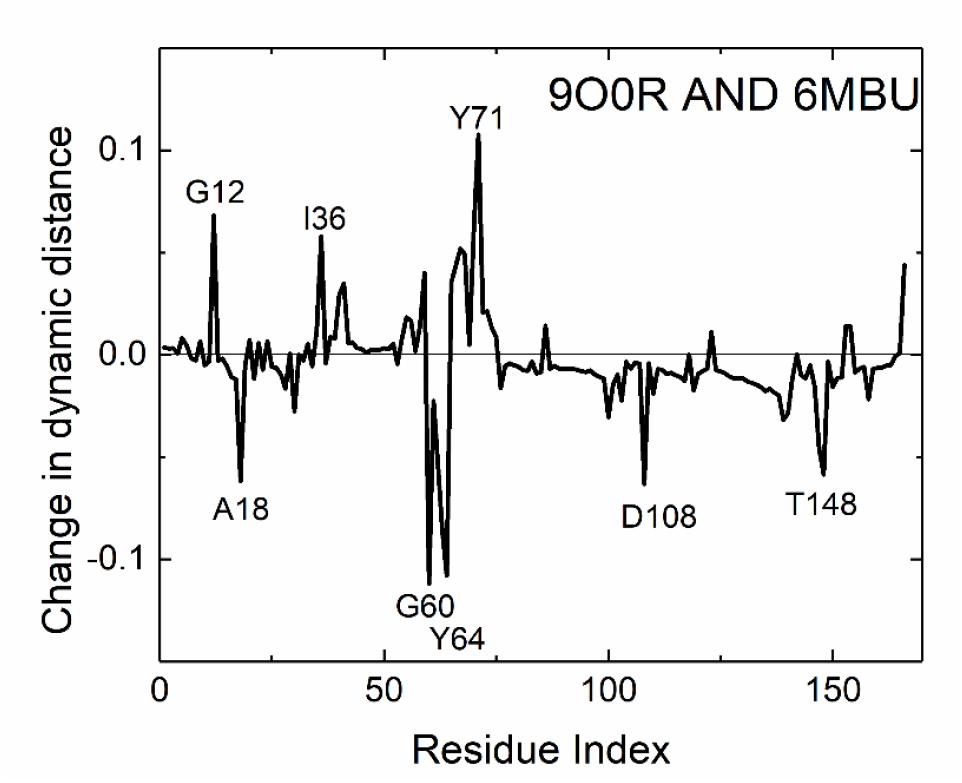
Change in dynamic distance patterns upon adagrasib binding to GDP-bound KRAS. Comparison between inhibitor-bound structure (9O0R.pdb) and apo GDP-bound KRAS (6MBU.pdb) using GNM with a cutoff radius of 7.8 Å. Dynamic distances were converted to z-scores within each structure to enable comparison. The ordinate shows Δz_12_,_j_ = z_12_,_j_(9O0R) - z_12_,_j_(6MBU) with G12 as reference residue. Values of *R*_*ij*_ are calculated from Eqs. 3-a and Eq. 6

**Figure 5.**
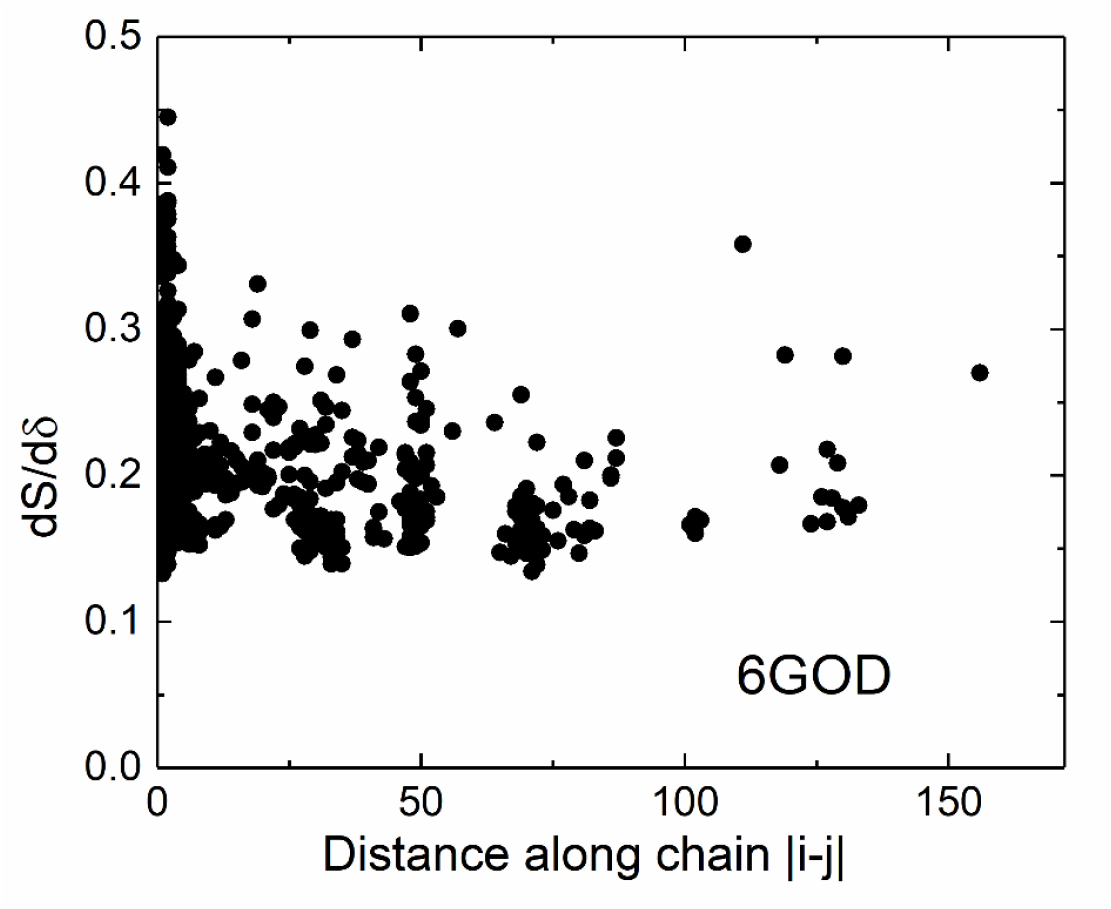
Increase in entropy for a unit decrease in edge weight, presented as a function of the distance between the two residues of the edge. The distance is taken as the absolute value of the difference between residue indices. Calculations are for the wild type protein 6GOD.pdb. Values of ∂*S* / ∂*δ* are calculated from Eq. 16.

Positive values indicate residue pairs whose dynamic distance increased upon inhibitor binding; these interactions become more critical bottlenecks as more spanning trees route through them, expanding the accessible configurational phase space. Negative values indicate decreased dynamic distance; these interactions become less critical, reducing their role in network communication. Peak positions are conserved across different choices of reference residue (data not shown), validating the robustness of these patterns.

### 3.3 Entropy sensitivity

Entropy is a global measure of conformational flexibility, while entropy sensitivity quantifies how this entropy changes when a specific residue–residue contact is perturbed. As expressed in 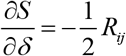, the change in entropy is directly determined by the *R*_*ij*_ of the perturbed edge: decreasing an edge’s centrality increases the protein’s entropy, and vice versa. In Figure 5, we show the entropy increase resulting from a unit decrease in edge weights for WT KRAS, plotted as a function of the sequence distance between contacting residues. Ignoring covalently bonded edges, the dependence on ∣i−j∣ is weak, indicating that long-range contacts contribute similarly to conformational flexibility as short-range contacts.

Our entropy-sensitivity analysis indicates a weak dependence of the entropic contribution on sequence separation. In the GNM-based metric used here, the mean entropy-change per contact shows little systematic decline with increasing ∣i−j∣, suggesting that at the level of coarse-grained conformational entropy both local and more distant contacts can carry comparable entropic weight. We emphasize two important caveats. First, this result concerns entropic contributions to global conformational flexibility computed from a Laplacian/pseudoinverse framework, not the detailed energetics or kinetic pathways of folding. Decades of folding work [50-52] demonstrate that local contacts frequently dominate early folding nucleation and enthalpic stabilization, properties determined primarily by energetic considerations. Our observation concerns a different quantity: the conformational entropy of the folded state as computed from network connectivity. In this context, we find that both local and long-range contacts contribute comparably to entropy per contact when assessed through the GNM Laplacian framework. These findings address different aspects of protein physics, folding energetics versus equilibrium conformational entropy, and are therefore not in conflict. Second, the GNM is a coarse-grained, isotropic model that neglects side-chain heterogeneity and detailed chemical energetics; therefore, our claim is limited to the architecture of entropy as assessed by network connectivity.

Having established the key residue interactions that serve as dynamic and topological bottlenecks in KRAS, we now examine their thermodynamic role from the point of view of entropy sensitivity. The direct relationship between entropy change and *R*_*ij*_ posits that the very edges critical for communication are also primary levers for global conformational entropy.

The high-edge-centrality contacts identified in Figure 1 correspond directly to regions of high entropy sensitivity. For instance, the E31-D30 switch I edge, which appears in 42% of all spanning trees, also possesses a high entropy sensitivity value of 0.21. This means that strengthening or weakening this single contact would have a disproportionately large effect on the overall configurational flexibility of KRAS. Similarly, the Q61-G12 edge, with its 33% spanning tree probability, is a significant thermodynamic control point. This is not a coincidence; it is a fundamental principle of the network: the contacts that are topologically indispensable for long-range communication are also the most efficient at modulating the system’s global flexibility.

This framework provides a thermodynamic interpretation for the network rewiring observed upon mutation. The G12D mutation, which significantly increases the dynamic distance *R*_*ij*_ for the E49-K172 pair, simultaneously increases its entropy sensitivity. This means the mutation not only makes this long-range conduit more topologically critical (increasing its spanning tree probability from 86% to 90%) but also amplifies its role as a thermodynamic gateway. Perturbations at E49 in the mutant will have a greater impact on global entropy than in the wild type, potentially altering the balance between active and inactive states and contributing to the constitutive signaling phenotype.

Conversely, interactions that lose dynamic distance upon mutation, such as those involving E63, see a corresponding reduction in their entropy sensitivity. This signifies a loss of thermodynamic influence, further marginalizing these pathways in the mutant’s functional landscape.

In summary, the entropy sensitivity metric shows that the allosteric control points of KRAS are integrated dynamic, topological, and thermodynamic hubs. Perturbations, whether pathological mutations or therapeutic ligands, exert their effects not only by rerouting information flow but also by directly reshaping the protein’s energy landscape, with the most profound changes occurring at these centrally located, high *R*_*ij*_ edges.

## 4. Discussion

This work introduces a unified theoretical framework for analyzing allosteric communication in proteins based on *dynamic distance R*_*ij*_, a dimensionless quantity derived from the graph-theoretical concept of *resistance distance* [33], which we interpret here as a measure of fluctuation differences between residues in the protein network. Three complementary measures emerge from this foundation: dynamic distance quantifies motion dissimilarity between residue pairs; edge centrality weights this by topological significance to capture spanning tree probabilities; and entropy sensitivity relates local perturbations to global conformational flexibility. The mathematical unity of this framework—all three measures being linear functions of *R*_*ij*_ reflects a fundamental principle: residue pairs that exhibit large relative fluctuations simultaneously serve as critical bottlenecks through which multiple communication pathways must pass, enabling the network to explore large configurational phase space while maintaining connectivity.

### Conceptual unification of fluctuations and network topology

The connection between mean-square relative fluctuations ⟨(Δ*R*_*i*_ − Δ*R*_*j*_)^2^⟩ and spanning tree probabilities may initially seem counterintuitive: why should poorly coordinated motion (high *R*_*ij*_) indicate communication importance? The resolution lies in recognizing that high *R*_*ij*_ edges function as flexible bottlenecks, connections through which many alternative communication pathways must pass to maintain network connectivity. A bottleneck in this context is an edge which is topologically essential, whose removal would eliminate numerous spanning trees, forcing the network to rely on fewer connectivity patterns. Conversely, an edge with low *R*_*ij*_ is redundant: alternative pathways can bypass it, so the network does not critically depend on that specific connection. Removing a high *R*_*ij*_ bottleneck would eliminate numerous alternative connectivity patterns and collapse the accessible configurational phase space. Thus, high dynamic distance identifies residue pairs that are simultaneously dynamically flexible and topologically critical, a combination characteristic of allosteric communication nodes that enable signal transmission between otherwise rigid functional domains.

### Biological interpretation requires functional context

Dynamic distance values alone do not determine biological significance. High *R*_*ij*_ can arise from functionally critical allosteric bottlenecks or from peripheral surface residues with high intrinsic flexibility. In KRAS, we observe that certain high *R*_*ij*_ pairs correspond to experimentally validated allosteric sites, strengthening their interpretation as critical communication nodes. Crucially, convergence across our three measures - dynamic distance, edge centrality, and entropy sensitivity - provides stronger evidence for functional importance than any single metric. This multi-metric validation helps distinguish genuinely important communication bottlenecks from incidental sources of motion dissimilarity.

### Perturbation analysis reveals network reorganization

Our KRAS analysis examines two representative perturbations: the oncogenic G12D mutation and adagrasib inhibitor binding. Changes in dynamic distance reflect network reorganization: increased *R*_*ij*_ indicates an edge becoming a more critical bottleneck as additional spanning trees route through it, while decreased *R*_*ij*_ indicates reduced criticality as alternative pathways emerge. The functional consequences depend on context. In wild-type protein, critical bottlenecks enable efficient allosteric signaling. For therapeutic inhibitors, reducing *R*_*ij*_ at these edges creates pathway redundancy that disrupts communication. For oncogenic mutations, increased *R*_*ij*_ may establish abnormal critical pathways driving constitutive signaling. The G12D mutation shows mixed effects-some edges become more critical while others become redundant - reflecting complex network rewiring. Adagrasib binding shows predominantly decreased *R*_*ij*_ values, consistent with disrupting critical communication bottlenecks through pathway redundancy.

### Understanding bottlenecks and redundancy

To interpret changes in dynamic distance upon perturbation, we must distinguish between bottleneck edges and redundant edges. A bottleneck is a connection that appears in many spanning trees; if removed, it would fragment the network or force communication through substantially fewer alternative pathways. High *R*_*ij*_ values identify such bottlenecks because they represent edges through which the network must route many of its connectivity patterns. In contrast, a redundant edge appears in few spanning trees because alternative pathways can easily bypass it; the network maintains connectivity even without that specific connection. When perturbations (mutation or ligand binding) increase *R*_*ij*_ for an edge, that connection becomes a more critical bottleneck-more spanning trees must route through it, concentrating communication pathways. When perturbations decrease *R*_*ij*_, the edge becomes more redundant-alternative pathways emerge that can bypass it, distributing communication across multiple routes. Neither outcome is inherently beneficial or detrimental; the functional consequence depends on whether efficient communication is desired (wild-type function requires bottlenecks) or should be disrupted (therapeutic inhibition benefits from creating redundancy).

### Limitations and scope

GNM’s isotropic assumption precludes analysis of directional coupling mechanisms. However, our focus is identifying which residue pairs exhibit strong dynamic coupling regardless of directionality. For allosteric networks, high *R*_*ij*_ identifies potential communication nodes whether coupling involves correlated, anticorrelated, or orthogonal motions. Validation against KRAS allosteric sites supports this position, suggesting isotropic analysis can identify functionally relevant patterns despite lacking directional information.

This study provides a proof of concept using KRAS as a single case study. Systematic benchmarking against other proteins, network methods, and experimental datasets remains important future work. While GNM provides convenient implementation, the theoretical framework applies equally to force-field-based or MD-derived networks, as *R*_*ij*_ reflects fundamental graph-theoretic properties independent of specific parameterization.

### Relationship to existing network analyses

Classical protein network analyses employ centralities (degree, betweenness, closeness, eigenvector) that emphasize local connectivity or geodesic paths [53-55]. Our measures extend these by explicitly linking theconceptof dynamic distance to global network properties through spanning trees and entropy. Previous work has examined related concepts - entropy changes upon binding [56], resistance distance as communication cost [57] - but not as a unified framework deriving three complementary perspectives from a single mathematical foundation. This integration represents the novel contribution of our approach.

### Spanning tree interpretation

Spanning trees are mathematical constructs, not literal physical pathways. They provide a rigorous statistical framework for quantifying how edges contribute to global connectivity. Edge participation probabilities reflect structural importance for network-level information flow. Our KRAS results show high-probability edges correspond to experimentally validated allosteric sites, supporting this statistical interpretation. However, these findings require further experimental validation to establish mechanistic pathways.

## Appendix A. Derivation of edge centrality

### 2.4.3 Spanning trees and probabilities

The probability of a spanning tree T is given by the contribution of that tree’s edge weights divided by the sum of the contributions of all spanning trees. Accordingly, the probability is

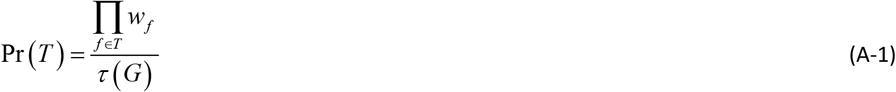

where the numerator is the product of the edge weights in T, and the denominator τ(G) is the sum of such products over all spanning trees.

This is like a Boltzmann probability in physics: heavier (larger weight) trees are more likely.

Probability of picking an edge: In the same way, we define the probability that a given edge e appears in a spanning tree drawn from this distribution:

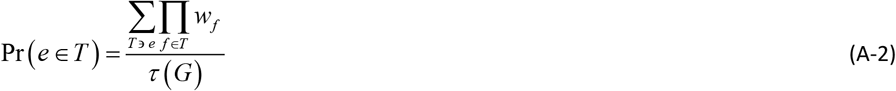

Here, the numerator is the sum of the weights of all spanning trees that contain e, and the denominator τ(G) is again the sum of such weights over all spanning trees. This gives a normalized probability between 0 and 1.

In Eq. A-2 we separate out *w*_*e*_ from the product as

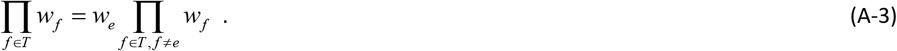

Then we differentiate with respect to 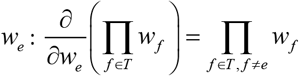. Summing over all the trees that contain e we obtain

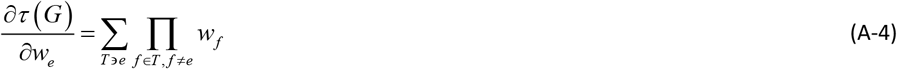

Using Eqs. A-3 and A-4 we obtain

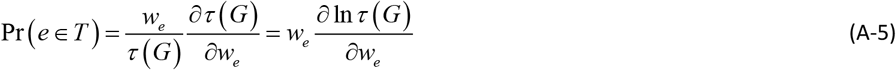

The logarithmic version of Kirchhoff’s matrix-tree theorem expressed in terms of eigenvalues is

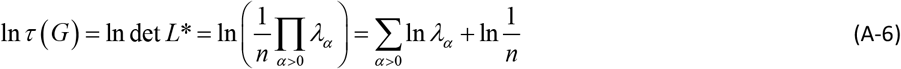

Differentiate with respect to *w*_*e*_

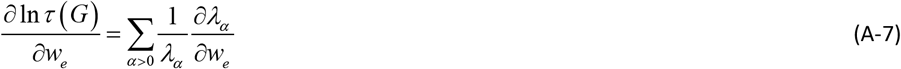

Denoting the eigenvetors of *L* by *u*_*α*_, eigenvalue perturbation gives

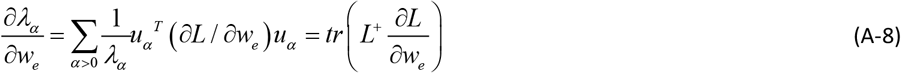

because 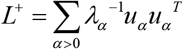 on the nonzero eigenspace.

The derivative ∂*L* / ∂*w*_*ij*_ may now be computed. For the edge *e* = (*i, j*)

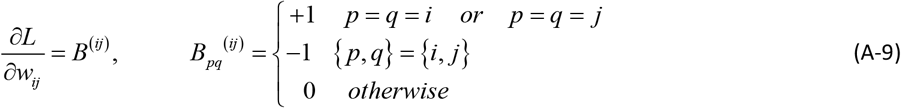

Therefore,

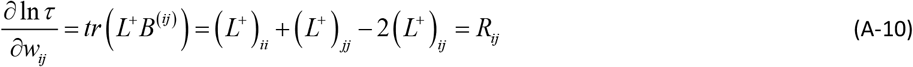

Substituting Eq. A-5 in Eq. A-10 we obtain

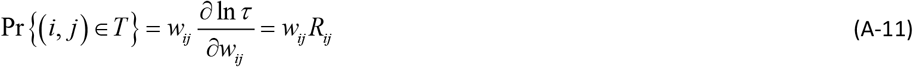

or

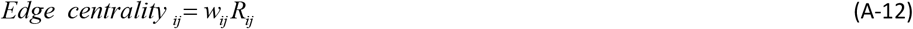

Thus, edge criticality quantifies the global importance of an inter-residue interaction: if *w*_*ij*_ = 1 it equals the fraction of spanning trees containing (i,j), and more generally it is the probability that the interaction participates in a randomly chosen weighted spanning tree, thereby capturing its role in inter-residue couplings at the global scale.

## Appendix B. Derivation of ∂*λ*_*k*_ / ∂*δ*^*+*^

Suppose the Laplacian depends smoothly on *δ* . Lt

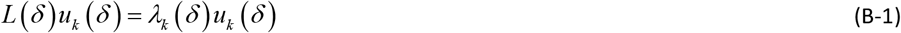

with eigenvectors normalized *u*_*k*_ (*δ*)^*T*^ *u*_*k*_(*δ*) = 1 . We want the derivative of *λ*_*k*_ (*δ*) with respect to *δ* . Differentiate both sides of the eigenvalue equation

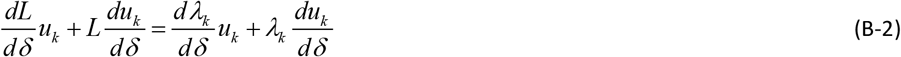

Now left-multiply by *u*_*k*_^*T*^ . Using orthonormality *u*_*k*_^*T*^ *u* = 1

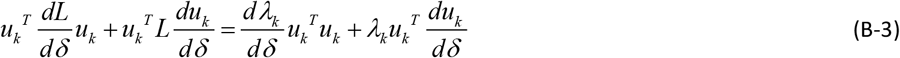

Simplify using 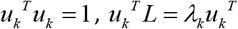, so

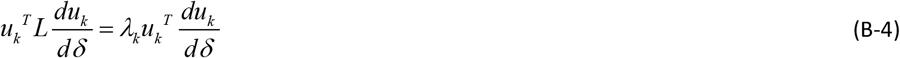

That cancels with the term on the right. What remains is the Rayleigh–Schrödinger expression:

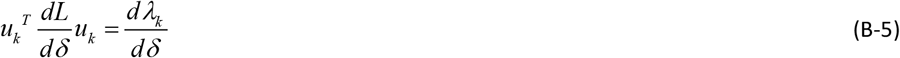

For the Laplacian, when we increase *w*_*ij*_ by *δ*, the perturbation is

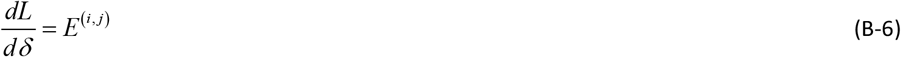

where, *E*^(*i, j*)^ is a sparse matrix with entries +1 at (i,i) and (j,j), −1 at (i,j) and (j,i), and 0 otherwise. Thus,

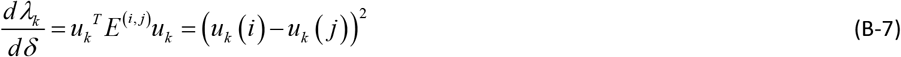

## Appendic C. Equivalence of Relative Fluctuations and Spanning-Tree Connectivity

From properties of the Laplacian and its pseudoinverse and Eq. A-10

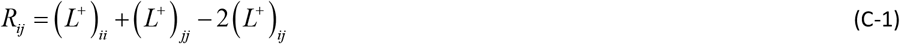

By the Matrix-Tree Theorem, the quadratic form given by Eq. C-1 is proportional to the fraction of spanning trees connecting residues i and j in the network.

From GNM

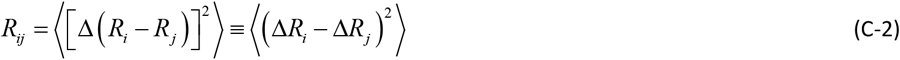

We can verbally state the equivalence of Eqs. A-1 and A-2 as:

The dynamic distance *R*_*ij*_ is defined from the Laplacian pseudoinverse and measures the mean-squared relative fluctuations between residues. By the Matrix-Tree Theorem, this same quantity is proportional to the fraction of spanning trees connecting the two residues. Thus, an increase in relative fluctuations between residues i and j is mathematically identical to an increase in the number of spanning trees connecting them.

## Notes

### Competing Interest Statement

The authors have declared no competing interest.

### Summary of Updates

The revised version introduces a unified mathematical framework based on three complementary measures of network organization derived from a single underlying quantity. The first, the dynamic distance Rij, quantifies the mean-squared relative fluctuation between residue pairs. From this foundation, two additional metrics are derived: the edge centrality, which identifies contacts critical for global connectivity by measuring their recurrence across all possible communication pathways, and the entropy sensitivity, which quantifies how perturbations to specific interactions alter system-wide flexibility. While these concepts were included in the original submission, they have been substantially clarified, corrected, and reformulated in the present version to improve mathematical and conceptual consistency.

